# Epigenetic vulnerabilities of leukemia harboring inactivating EZH2 mutations

**DOI:** 10.1101/2023.11.20.567858

**Authors:** Mona A. Alqazzaz, Genna M. Luciani, Victoria Vu, Raquel Martinez Machado, Magdalena M. Szewczyk, Ella C. Adamson, Sehyun Cheon, Fengling Li, Cheryl H. Arrowsmith, Mark D. Minden, Dalia Barsyte-Lovejoy

**Affiliations:** Structural Genomics Consortium, University of Toronto, Toronto, Ontario, Canada; Princess Margaret Cancer Centre, Toronto, Ontario, Canada; Department of Medical Biophysics, University of Toronto, Ontario, Canada; Department of Pharmacology and Toxicology, University of Toronto, Ontario, Canada

**Keywords:** Epigenetics, EZH2, Histone methylation, Acute myeloid leukemia

## Abstract

Epigenetic regulators such as the polycomb repressive complex 2 (PRC2) play a critical role in both normal development and carcinogenesis. Mutations and functional dysregulation of PRC2 complex components such as EZH2 are implicated in various forms of cancer and associated with poor prognosis. This study investigated the epigenetic vulnerabilities of acute myeloid leukemia (AML) and myelodysplastic/myeloproliferative disorders (MDS/MPN) by performing a chemical probe screen in patient cells. Paradoxically, we observed increased sensitivity to EZH2 and EED inhibitors in AML and MDS/MPN patient cells harboring *EZH2* mutations. Expression analysis indicated that EZH2 inhibition elicited upregulation of pathways responsible for cell death and growth arrest, specifically in patient cells with mutant EZH2. The identified *EZH2* mutations had drastically reduced catalytic activity, resulting in lower cellular H3K27me3 levels and were associated with decreased EZH2 and PRC2 component EED protein levels. Overall, this study provides an important understanding of the role of EZH2 dysregulation in blood cancers and may indicate disease etiology for these poor prognosis AML and MDS/MPN cases.

## I. Introduction

Epigenetic regulation encompasses the covalent modification of histone proteins, such as acetylation, phosphorylation, and ubiquitylation, resulting in the activation or inhibition of gene transcription and regulation of chromatin structure (1). Dysregulation of histone methylation is associated with various stages of carcinogenesis, including tumor initiation, promotion, and progression. Thus, targeting histone methyltransferases is being explored as a cancer therapeutic approach (2). The polycomb repressive complex 2 (PRC2), which is responsible for the mono-, di- and tri-methylation of histone 3 at lysine 27 promotes the compaction of the nucleosome and the repression of genes involved in development, cell differentiation, and proliferation (3).

The core subunits that form human PRC2 are the enhancer of zeste homologs 1 or 2 (EZH1 and EZH2), embryonic ectoderm development (EED), suppressor of zeste 12 (SUZ12), and retinoblastoma suppressor associated protein 46/48 (RbAp46/48). Multiple auxiliary subunits also enhance PRC2 chromatin association and enzymatic activity (3). EZH2 (or EZH1) is the primary catalytic component of PRC2, which is mainly mediated by its SET and CXC domains at its C-terminus (4). EED plays a regulatory role by stimulating EZH2/EZH1 activity, SUZ12 docks the PRC2 complex to the nucleosome, while other subunits regulate the enzymatic activity and genome localization of the PRC2 complex (5,6).

Numerous studies have implicated dysregulated EZH2 function in carcinogenesis (7,8). Depending on the type and stage of cancer, EZH2 has been shown to promote or repress cancer (9,10). For instance, EZH2 overexpression is observed in various solid tumors e.g. prostate, bladder, and breast cancers, and recurrent gain of function mutations of EZH2 have been identified in diffuse large B cell lymphoma and non-Hodgkin’s lymphomas suggesting that EZH2 acts as an oncogene (7,8). Additionally, loss of function mutations in epigenetic regulators, such as lysine demethylase 6A (UTX) and SWI/SNF components, that functionally oppose PRC2, have been identified in multiple cancers and shown to have synthetic lethal relationships with EZH2 (11,12). Taken together, these data provide strong evidence for a pro-oncogenic function by EZH2; currently, the EZH2 inhibitor tazemetostat is approved for clinical use in lymphomas harboring EZH2 activating mutations and in SWI/SNF mutant epithelioid sarcoma (8,13).

In contrast, myeloid malignancies present with inactivating *EZH2* mutations at diagnosis or during disease progression and relapse, most common in MDS/MPN (8-13%), MDS (3-13%) (14,15) and in 1% of de novo AML patients (16,17). These loss of function (LOF) *EZH2* mutations are correlated with lower survival rates (14,18-25). In addition to LOF mutations, splice site mutations and splicing factor mutations have been reported to disrupt EZH2 in hematological malignancies (26,27). Interestingly, germline inactivating/LOF mutations in PRC2 components, EZH2, SUZ12 or EED, are found in patients with overgrowth syndromes, Weaver syndrome, and Weaver-like syndrome and a subset of these patients have an increased risk of developing myeloproliferative neoplasms (14,28). The dual tumor suppressor and oncogene nature of EZH2 is illustrated by disease models of leukemia, where EZH2 suppressed leukemogenesis while facilitating the disease maintenance cancer (9,10).

In this study, we performed an epigenetic vulnerability screen in AML/MDS patient cells. Surprisingly, *EZH2* mutations (*EZH2^MUT^*) in AML/MDS rendered cells more sensitive to PRC2 catalytic function inhibition. These cells have significantly reduced baseline levels of H3K27 methylation and respond to further PRC2 inhibition by regulating the pathways responsible for cell death and growth arrest. The novel *EZH2* mutations we identified not only resulted in reduced H3K27me3 but also lower levels of EZH2 protein and, in some cases, destabilization of the PRC2 complex. Our study provides an important understanding of the role of EZH2 dysregulation in blood cancers.

## II. Methods

### 1. AML patient cell culture and viability assessment

Patient cells were collected with informed consent in accordance with the procedures outlined by the University Health Network’s Research Ethics Board approval. Cryopreserved, ficoll purified patient mononuclear cells from either bone marrow or peripheral blood that had been stored in liquid nitrogen were rapidly thawed, washed and cultured as described before (29), details in the supplementary material. Patient cells were transferred onto OP9 cells in 96-well plates for compound testing at a density of either 25000 or 50000 cells per well in 100 μL media with 0.1% DMSO/compound (compound concentrations used in Suppl Table 1). Viable cell number was assessed at day 12-14. To assess viability, we used Sytox Blue (Life Technologies) staining and flow (Supplementary methods).

### 2. H3K27me3 and EZH2 flow cytometry and immunoblotting

H3K27me3 methylation and EZH2 levels in patient cells were determined using flow cytometry as previously described (30). Briefly, cells were collected, fixed, permeabilized and stained with H3K27me3 or EZH2 antibodies with OCI-AML-20 cells (29) treated with and without EPZ-6438 were used as a control for H3K27me3 staining. Full methods and staining validation data are provided supplementary materials and Suppl Fig 1.Patient cells were lysed, protein harvested, and western blots were run as previously described (31). Blots were incubated overnight at 4°C with the antibodies as described in Supplementary Table 2.

### 3. Histone methyltransferase assays

EZH2 Wild type (*EZH2^WT^*) and mutant (*EZH2^MUT^*) and other components of PRC2 complex were expressed and purified and H3K27 methylation assays was performed as we previously described (32) see supplementary methods for details. In brief, EZH2 trimeric or pentameric complex were used and the incorporation of a tritium-labeled methyl group to lysine 27 of H3 (21–44) peptide was monitored using a scintillation proximity assay (SPA). Reactions were performed in triplicate, and IC_50_ values were determined by fitting the data to a four-parameter logistic equation using GraphPad Prism 8 software.

### 4. RNA sequencing

*EZH2^WT^* or *EZH2^MUT^* patient cells were treated with DMSO or 1 μM EPZ-6438 for four days. Each condition included three biological replicates. RNA purification, library preparation for sequencing, and detailed sequence processing and subsequent analyses can be found in the Supplementary methods.

## III. Results

### 1. Epigenetic chemical probe screen identifies that leukemia patient cells harboring EZH2 mutations are sensitive to EZH2 inhibitor

We screened a library of 25 well-characterized epigenetic chemical probes for suppression of proliferation of 21 primary AML samples; cells were from patients at presentation or relapse. The top panel of Figure 1 shows a mutational background of the AML patient cells.

**Figure 1:**
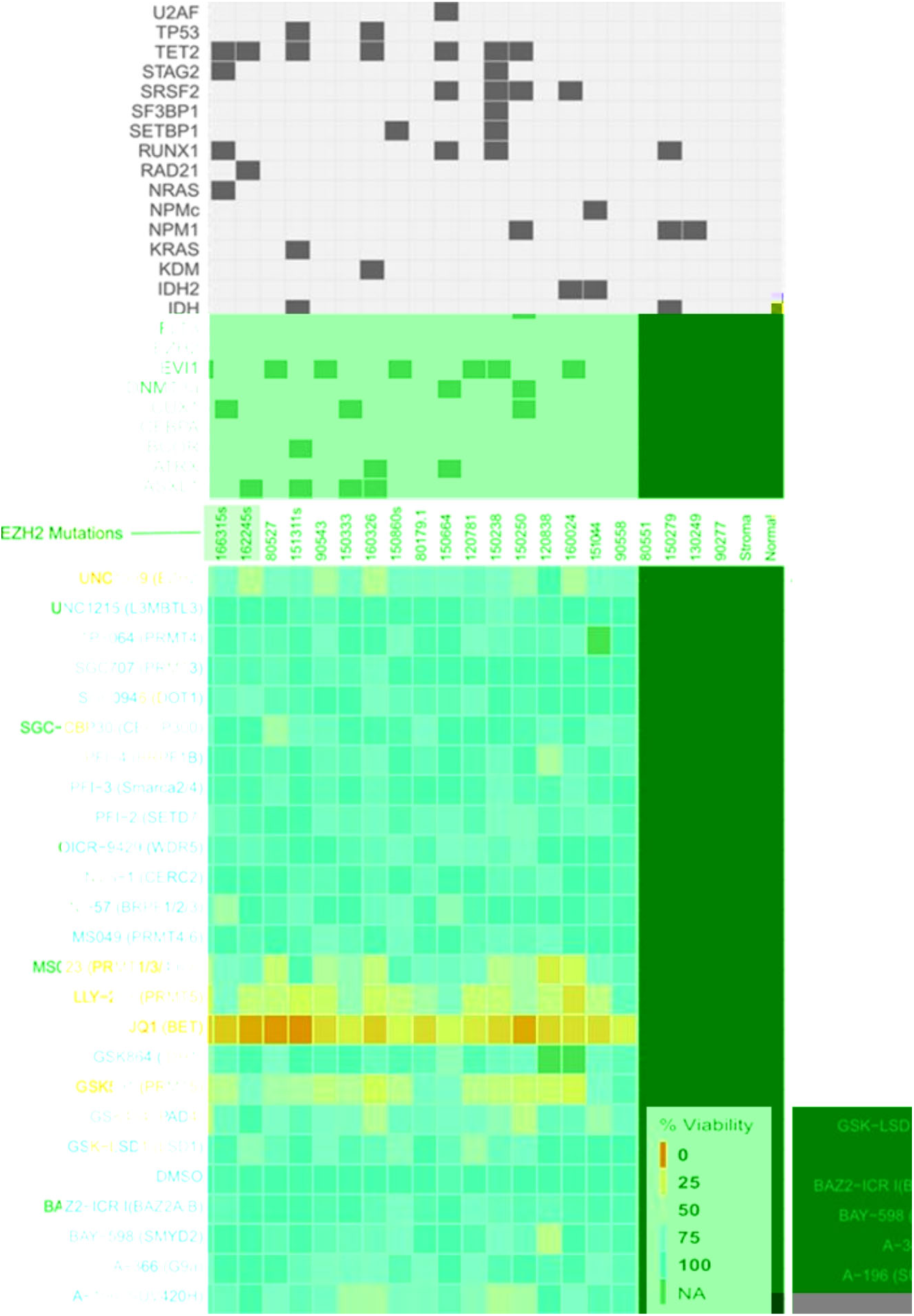
Epigenetic chemical probe screen identifies AML samples hypersensitive to EZH2 inhibition. AML sample mutation data is shown at the top. Bottom heatmap: Viable cell count heatmap of epigenetic chemical probe screen in leukemia patient primary cells. The screen was performed on 21 patient primary AML/MDS cells, one control normal PBMC sample and stroma cells used to support the AML cultures. Cells were grown for 12–14 days in the presence of compounds at concentrations indicated (Supplementary Table 1). Data are represented as mean viability calculated as a percentage of DMSO control (n = 4). Dark red indicate a decrease in cell viability relative to vehicle (DMSO). Grey squares indicate probe and sample combinations that were not screened.

The chemical probe library was selected to encompass small-molecule selective inhibitors of methyltransferases, bromodomains, acetyltransferases, and deacetylases. Patient cells, healthy donor peripheral blood mononuclear cells (PBMC), and stroma cells (used for maintenance of the patient cells) were treated for 12–14 days, and cell viability was measured using flow cytometry. JQ1, a bromodomain inhibitor, demonstrated uniform antiproliferative effects that are consistent with reports on many cancer cell lines (33). The EZH2 probe, UNC1999, potently inhibited cell proliferation of only two leukemia samples which surprisingly harbored EHZ2 mutation(s) (bottom panel of Figure 1). Since UNC1999 demonstrated increased potency in patient cells with *EZH2^MUT^*relative to *EZH2^WT^*, we proceeded to validate findings in a larger cohort of primary patient cells with EZH2 mutations.

### 2. Characterization of EZH2 mutations in AML patient cells

Despite the rarity of EZH2 mutations in leukemia, we obtained an additional six cryopreserved *EZH2^MUT^* leukemia patient samples from the Princess Margaret BioBank. We also used four leukemia patient cell samples with no *EZH2* mutations as controls for our experiments (Table 1, Figures 2A, 2B). Patient cells 177391 (Q653K), 170396 (V679M), and 166315 (E137X*term) harbored single *EZH2* mutations with an allele frequency of 91- 95.5% (Table 1). Patient cells 161821 (C576Y, R690H), 165009 (P527H, C552R), and 162245 (T683I, C538Y) had two mutations in *EZH2* with an allele frequency of (42-49%) each. These missense mutations were found in the catalytic SET and CXC domains required for the catalytic function of *EZH2* (mutations are highlighted in red in Figure 2A, B). The most commonly co-occurring mutations in *EZH2^MUT^*patient cells involved *ASXL1* and *TET2* (Figure 2C), consistent with previous findings (20,26,34).

**Figure 2:**
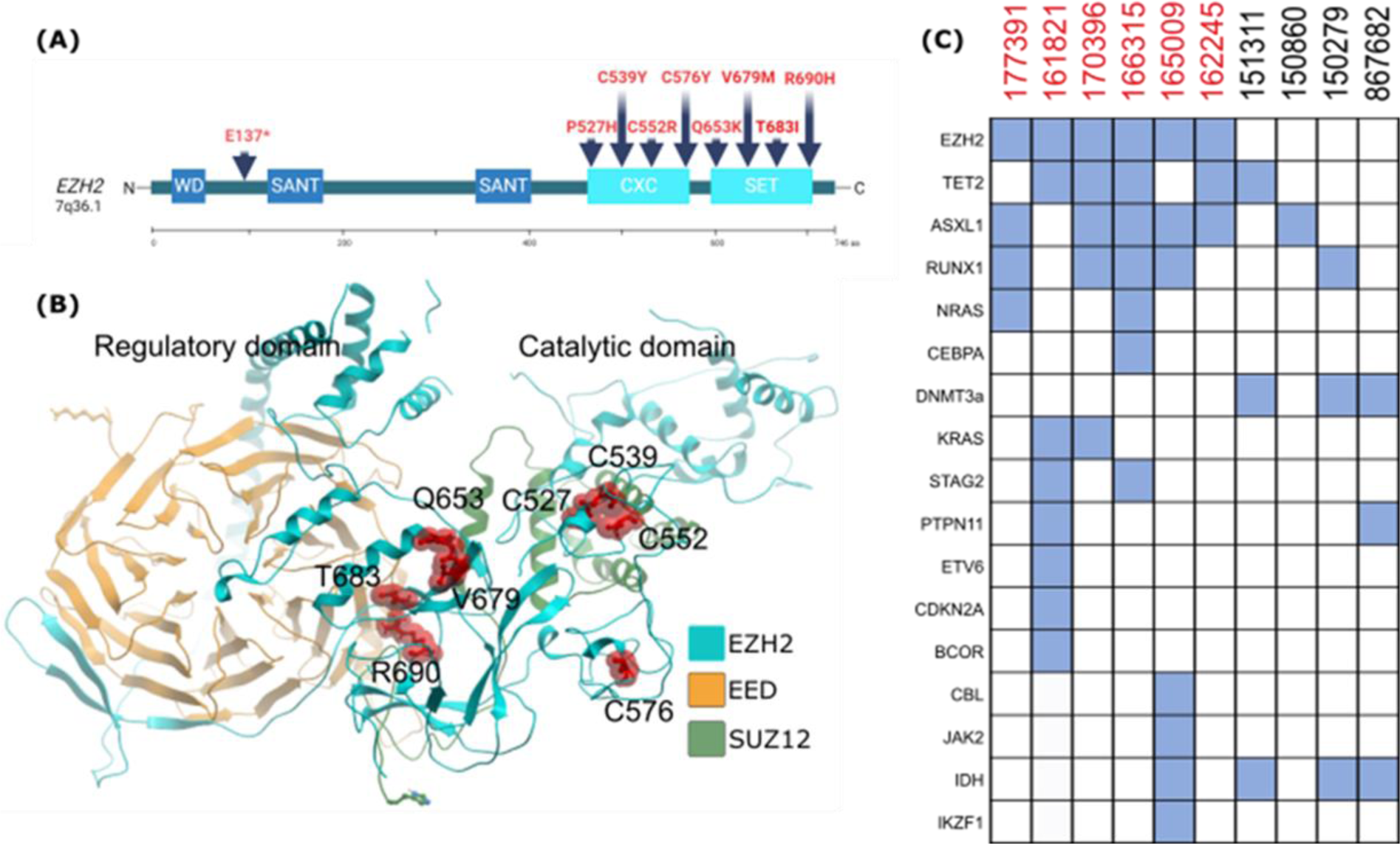
Summary of the mutation positions in EZH2 protein for the AML patient cells used. (A) Linear representation of EZH2 domains and the localization of mutations. Most mutations are located on the CXC and SET domains. (B) cartoon protein representation showing the localization of the missense mutations as amino acid residues (red sticks) in the protein structure (PDB ID 5HYN). (C) Summary of the main co-existing mutations in patient samples.

**Table 1:**
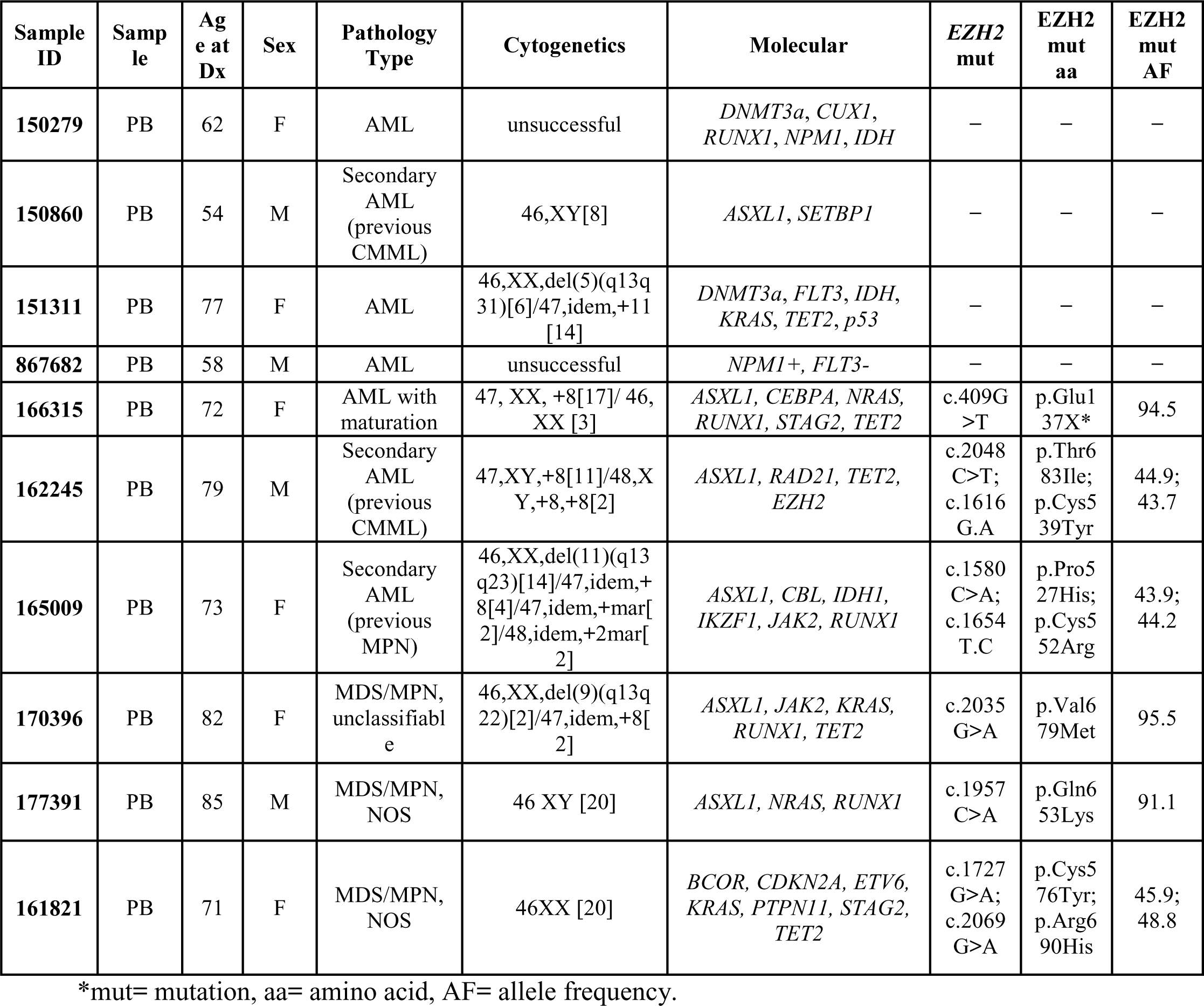
summary of EZH2 mutations and their allele frequencies in AML patients.

### 3. *EZH2^MUT^* patient cells have an increased sensitivity to PRC2 inhibitors relative to *EZH2^WT^*

To validate our initial AML patient screen findings, we determined the sensitivity of *EZH2^WT^* and *EZH2^MUT^* -patient cells to clinical EZH2 inhibitor EPZ-6438 (Tazemetostat, Tazverk). In this analysis, we also included a comparison to the EED inhibitor (A395) because of its ability to allosterically inhibit PRC2 at its regulatory peptide-binding site, thus providing the potential to inhibit both EZH2 and EZH1 (35). Cells were maintained in a co-culture system with OP9 stromal cells and growth factors treated with inhibitors for 14 days, at which time viable cells were analyzed using flow cytometry. We observed potent growth suppression in *EZH2^MUT^* AML/MDS primary patient cells with EPZ-6438 and A395, while growth suppression of other non-EZH2 mutant AML patient cells required much higher compound concentrations (Figure 3). Patient cells with *EZH2^MUT^* were hypersensitive (demonstrating lower half maximal inhibitory concentration (IC_50_)) to EPZ-6438 (0.03-0.4 µM) and A395 (0.02- 0.03 µM) inhibitor treatment compared to cells with *EZH2^WT^* (EPZ-6438 IC_50_ range=1.5-2.4 µM and A395 IC_50_ range = 1.2-2.9 µM) (Figure 3A, B). In contrast, the inactive (negative control) compound for the EED inhibitor (A395N) had no effect on the viability of any of the cells, indicating the inhibitors are targeting specifically the PRC2 activity (Figure 3A right graph). *EZH2^MUT^*patient cell response to A395 is illustrated by a left shift of the concentration-response curves (Figure 3C).

**Figure 3:**
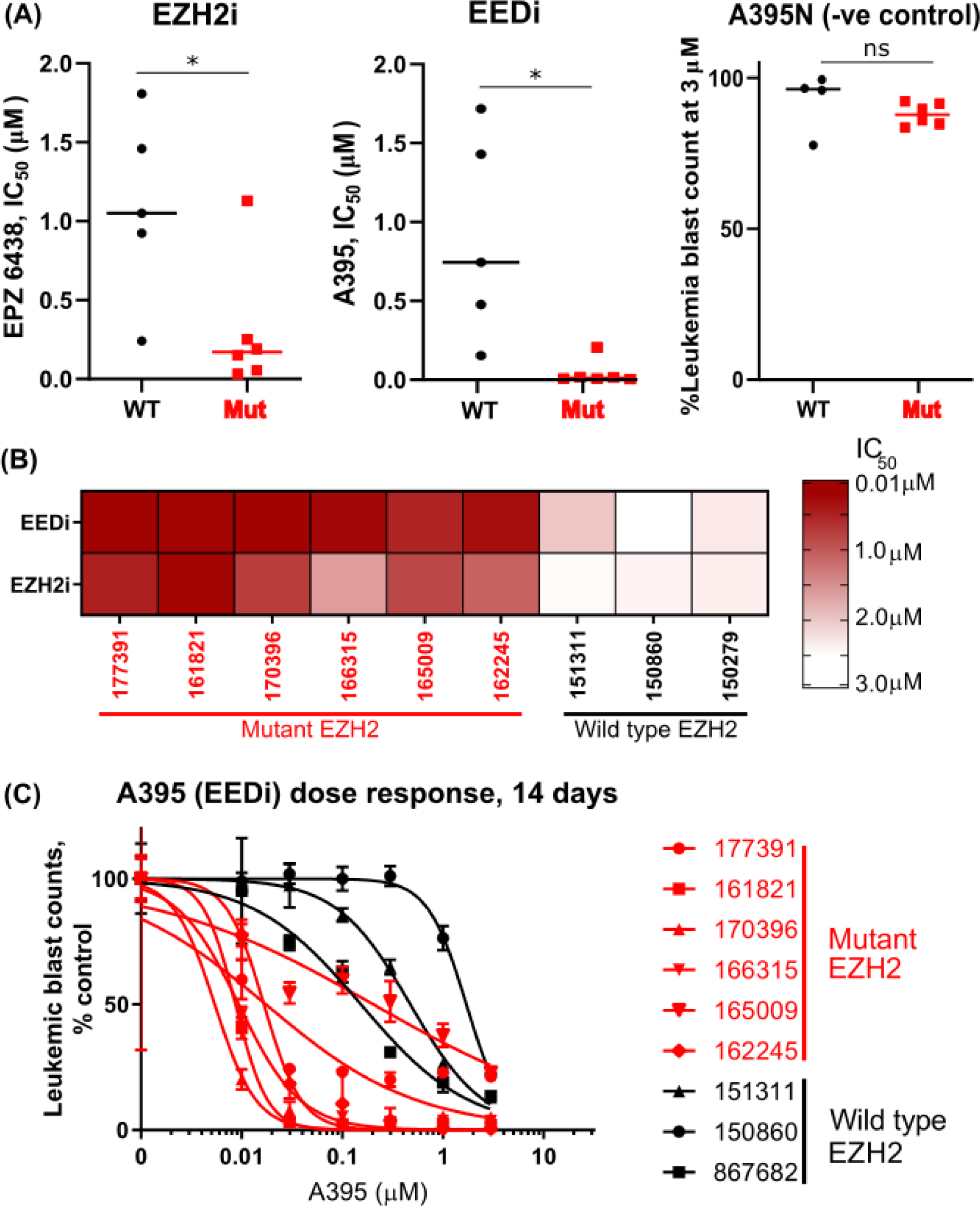
Validation of the epigenetic chemical screen using a larger number of AML patient cells with and without EZH2 mutations. (A) EZH2 Mutants are hypersensitive to EZH2 and EED Inhibition. Comparison of IC_50_ of patient samples with mutant and wildtype EZH2 using EZH2 inhibitor, EPZ 6438, and EED inhibitor, A395 and negative control A395N. IC_50_ was measured using flow cytometry monitoring leukemia cell blast count at 14 days of treatment with EZH2 or EED inhibitor. Comparisons were made using unpaired T-test, p= 0.002, 0.0041 and 0.2045, respectively. (B) heatmap representation of the results showing higher potency (red) vs lower potency (white). (C) Representative dose-response curves used to determine the IC_50_ values.

### 4. Characterization of EZH1, EZH2, and H3K27me3 protein levels in patient cells

Next, to determine if EZH2 mutations significantly affected H3K27 methylation, we determined trimethylation of lysine 27 of histone 3 (H3K27me3) mark using immunoblotting (Figure 4A). Patient cells with *EZH2^MUT^* had lower H3K27me3 mark levels than *EZH2^WT^* samples. To support the results of immunoblotting, we investigated the H3K27me3 mark levels using flow cytometry. We performed flow cytometry of CD45-positive AML patient cells and demonstrated that *EZH2^MUT^* - cells have significantly lower levels of H3K27me3 compared to *EZH2^WT^* - cells (Figure 4B).

**Figure 4:**
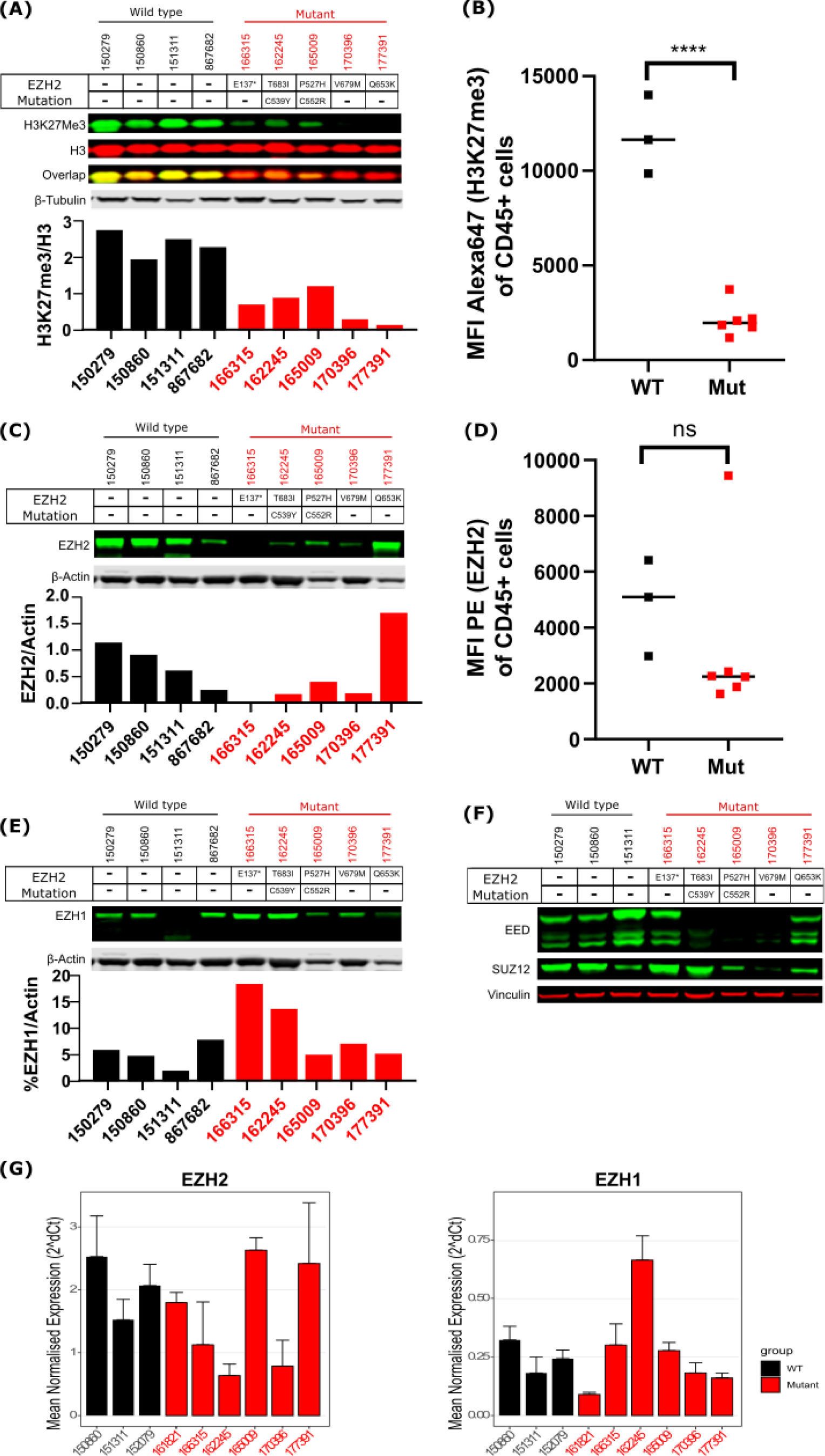
Characterization of PRC2 complex components and H3K27me3 levels in *EZH2^MUT^* and *EZH2^WT^* cells. (A) Western blot analysis demonstrating that the levels of H3K27me3 are significantly reduced in *EZH2^MUT^* patient cells (B) Flow cytometry also showed reduced levels of H3K27me3. P< 0.0001 using unpaired t-test (C) Western blot analysis of EZH2 levels in patient cells. Lower EZH2 levels were detected in most *EZH2^MUT^* patient samples. The bottom panel shows the quantification of EZH2 levels relative to actin. (D) Flow cytometry showed reduced levels of EZH2 in five out of six patient samples containing *EZH2^MUT^* in comparison to the wild type. Sample 177391 had high levels of EZH2 when measured both using western blot and flow cytometry. (E) Western blot analysis demonstrating levels EZH1 in patient cells with *EZH2^MUT^*and *EZH2^WT^*. (F)Western blot analysis of PRC2 components EED and SUZ12 (G) qPCR analysis of mRNA levels of EZH2 and EZH1 in patient samples. n=4 technical replicates. Data = mean ± SD.

We further determined the relative protein levels of EZH2 using immunoblotting (Figure 4C). Remarkably, EZH2 levels were lower in *EZH2^MUT^*- patient cells except for 177391, which had higher EZH2 levels. To further validate the immunoblotting results, we performed flow cytometry of cells cultured for two days and analyzed EZH2 levels in leukemic cells (CD45 positive). Consistent with immunoblotting data, *EZH2^MUT^* cells from the majority of patients had detectable but lower levels of EZH2 protein relative to *EZH2^WT^* cells (Figure 4D).

We also measured EZH1 protein levels and found it was variable across *EZH2^MUT^* and *EZH2^WT^* patient cells using western blot analysis (Figure 4E). *EZH2^MUT^*-162245 and 166315 had higher EZH1 levels, while the remaining three samples had an EZH1 protein range comparable to the *EZH2^WT^* samples. Interestingly, sample 166315 with E137X*term nonsense mutation had nondetectable EZH2, high levels of EZH1, and low levels of H3K27me3.

As the lower levels of EZH2 found in the *EZH2^MUT^* patient samples could potentially result in the destabilization of the PRC2 complex, we investigated the levels of EED and SUZ12. Interestingly, three out of five *EZH2^MUT^*samples had lower levels of EED, and two had low levels of SUZ12 (Figure 4F). We also compared the mRNA levels of *EZH2* and *EZH1* using qPCR. No significant trends were noted in mRNA levels between patient cells for both *EZH2^WT^* and *EZH2^MUT^* samples (Figure 4G).

### 5. EZH2 dependent gene expression program analysis

To gain insight into the mechanism underlying the observed increase in sensitivity of *EZH2^MUT^* patient cells, we performed RNA sequencing analysis (RNA-Seq) for two *EZH2^WT^* and two *EZH2^MUT^* patient samples. We treated the cells for four days with 1 µM EPZ-6438 or DMSO, selecting this treatment time because H3K27me3 levels were downregulated, but no significant cell death was observed at this time point. RNA sequencing analysis showed that EPZ-6438 differentially regulated 284 genes in *EZH2^WT^* -151311, 718 genes in *EZH2^WT^* -150860, 926 genes in *EZH2^MUT^*-155009 and 477 genes in *EZH2^MUT^* -162245 (Figure 5A).

**Figure 5:**
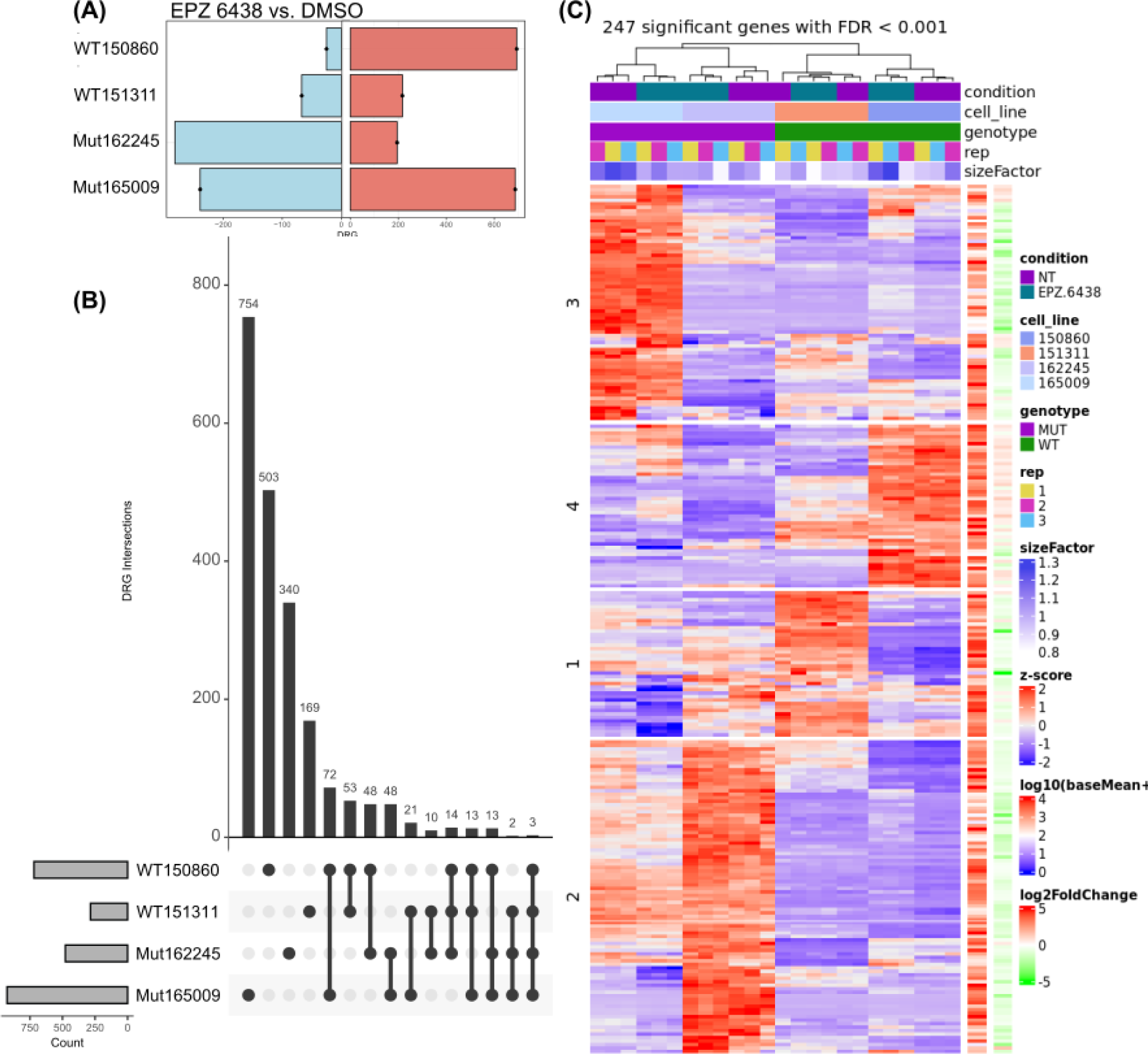
Summary of gene expression changes in response to EZH2 inhibition. (A) EPZ-6438 upregulated and downregulated differentially regulated genes (DRGs) in ***EZH2^WT^*** and ***EZH2^MUT^*** patient cells. (B) Comparison of DRGs between groups. The Horizontal Bar graph (blue) indicates the total number of DRGs in each patient cell upon treatment with EPZ-6438 vs DMSO. The circles in the matrix denote the number of unique DRGs in the sample. Connecting lines between circles indicates the overlap between groups. The vertical bar graph shows the number of unique or overlap datasets in the groups indicated in the matrix below. (C) Heatmap depicting clustering analysis of gene regulation by EPZ-6438. Patient gene expression is annotated with mutational status and treatment. The heatmap shows the log2 fold changes of the genes as colors, ranging from blue (negative) to white (zero) to red (positive).

We used an Upset plot to summarize the overlap of differentially regulated genes (DRGs) in *EZH2^MUT^* and *EZH2^WT^* patient cells treated with EPZ-6438 vs DMSO. In Figure 5B, the bottom left grey bar graph shows the total number of DRGs (up and down-regulated). The circles in the matrix panel represent unique or overlapping DRGs, with connected circles indicating overlap between datasets. This plot reveals minimal overlap of DRGs across all four datasets, with the majority being unique to each patient’s cells. The three genes commonly regulated in all four datasets are *CDHR1* (calcium-dependent cell adhesion molecules), *PTPRF* (tyrosine phosphatase involved in cell division and differentiation), and *SLC22A31* (predicted transmembrane transporter activity). K-means clustering analysis revealed that *EZH2^WT^* and *EZH2^MUT^* samples were more similar within their groups (see dendrogram at the top of Figure 5C), and the EPZ-6438 treatment responses were distinctly characteristic within samples, suggesting that there is an appreciable broad change in expression patterns in response to drug treatment when compared to control DMSO condition (Figure 5C).

Individual analysis of the *EZH2^WT^* sample response to EPZ-6438 indicated that genes associated with apical junction, estrogen response (*HES1*), and epithelial-mesenchymal transition (EMT) (*FBLN1, TGFBR3, GREB1, and ITGB5*) were upregulated (Figure 6, Supplementary Figure 2). The known *EZH2* target genes *TGFBR3, GREB1*, and *ITGB5* were upregulated (ENCODE). As reported before, the upregulation of LIN28B was also noted in the case of *EZH2^MUT^* sample 162245 (9). This is consistent with the repressive function of PRC2 on these genes. Overall, in *EZH2^WT^*samples, the top categories of EPZ-6438 regulated gene terms were associated with EMT and inflammatory response, while in *EZH2^MUT^* sample cells, predominant regulated pathways were: inflammation and cell cycle associated (Figure 6). Interestingly *EZH2^MUT^* cells responded to EPZ-6483 by prominently downregulating genes associated with inflammatory response (*IRF4*), survival and growth factor (*IL6*), DNA repair *(RAD51, POLD2*), proliferation E2F (*RRM2*), and MYC (*PLK4*) (Figure 6, Supplementary Figure 2). Concurrently, the upregulated genes were associated with the P53 pathway and growth arrest (CTSF, CDKN2A). Thus, despite very different genetic backgrounds, common between both *EZH2^MUT^* patient cells, EZH2 inhibition elicited the upregulation of genes associated with cell death and growth suppression pathways, while the genes responsible for proliferation and DNA repair were downregulated.

**Figure 6:**
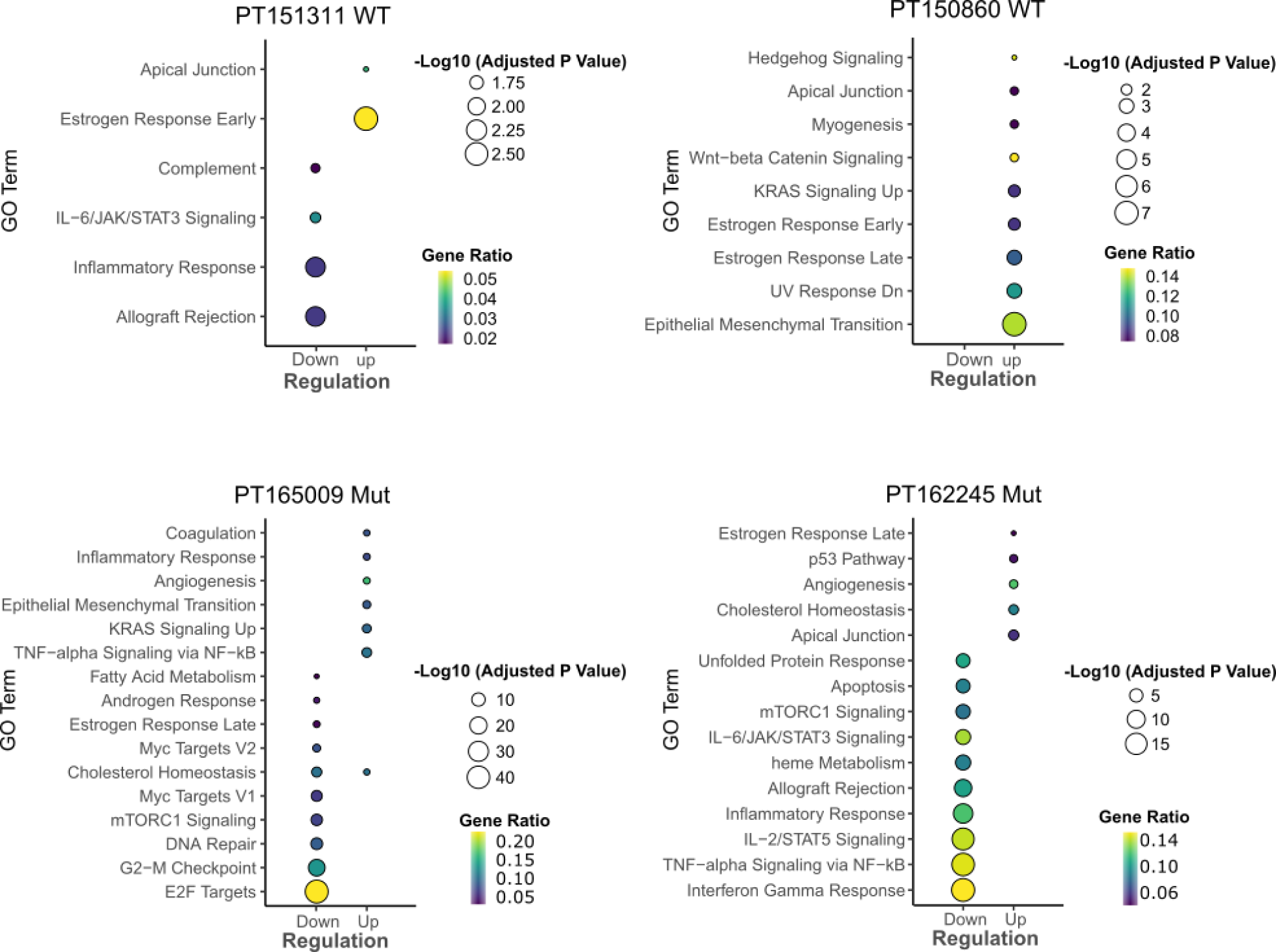
Gene Ontology (GO) Term Analysis for *EZH2^WT^* (top panel) and *EZH2^MUT^* (bottom panel). The graphs represent and individual analysis of two samples *EZH2^WT^* patient (151311 and 150860) and two *EZH2^MUT^* patient (162245 and 165009) cells. Differentially expressed genes are also shown as volcano plots (Supplementary Figure 2). Significantly (p<0.05) regulated biological terms from MSigDB represented as dot blot graphs with genes regulated out of total in the gene category indicated.

### 6. *EZH2^MUT^* have lower histone methyltransferase activity

As the *EZH2* mutations resulted in lower levels of the protein (Figure 4), it is possible that the observed decrease in H3K27me3 is due to lower EZH2 protein levels. Thus, it was unclear if the *EZH2* mutations in primary patient cells indeed cause a loss of function. To address this, we measured the *in vitro* methyltransferase activity of individual mutations in recombinant EZH2 using a scintillation proximity assay. The *EZH2* CXC domain mutations C576Y, C552R and SET domain mutations T683I, R690H, and V679M resulted in a complete loss of function (Figure 7A). EZH2-P527H retained a significant amount of PRC2 methylation activity, whereas EZH2-C539Y retained partial methylation activity (25%) in comparison to *EZH2^WT^* (Figure 7A).

**Figure 7:**
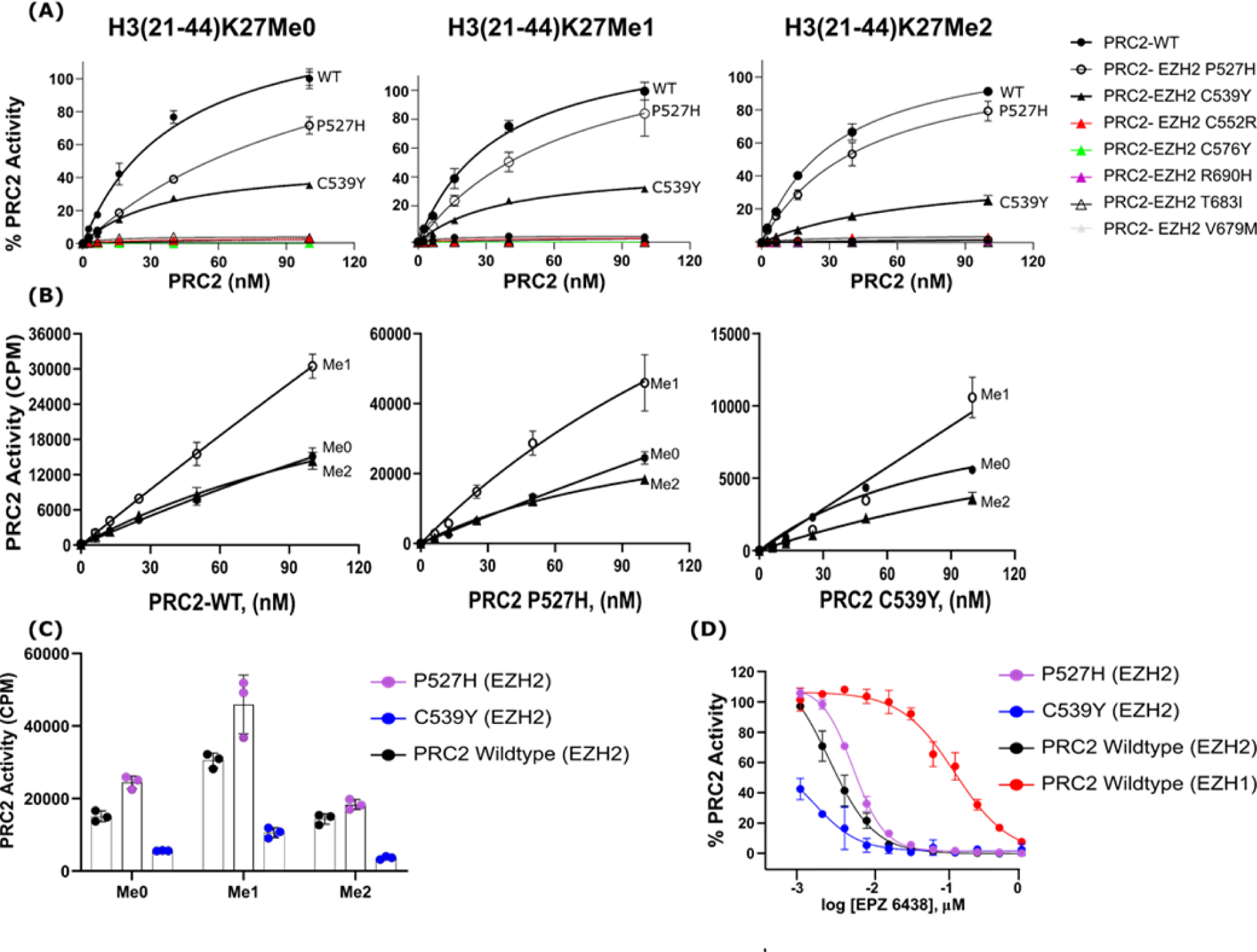
*In vitro* characterization of EZH2 mutations for the enzymatic activity using 3H-SAM incorporation methylation assay. (A) The catalytic activity of purified EZH2^WT^, and EZH2^MUT^ containing single mutations from AML patients was determined using unmethylated peptide to test overall methylation activity, mono and di- methylated H3K27 2- 44 biotinylated peptides to test for mutant methylation activity of various substrate using tritium labeled SAM. (B) a comparison of the *in vitro* substrate preference of EZH2^WT^ and P527H and C539Y mutant EZH2. All exhibited preferred activity towards the monomethylated histone peptide relative to the dimethylated and unmethylated peptides. (C) A comparison of the maximum PRC2 activity of EZH2^WT^ (black) and EZH2-C539Y (blue) and EZH2-P527H (purple). The maximum activity is lower for EZH2-C539Y but unchanged for EZH2-P527H for all three substrates. (D) EZH2-C539Y (blue) and EZH2-P527H (purple) sensitivity to EPZ-6438 is unchanged relative to EZH2^WT^ -containing PRC2. Experiments were performed in triplicates. Data= mean ± SD.

We then compared the methylation activity using mono-, and di- methylated peptide substrates to unmethylated peptides to determine if mutations in *EZH2* caused a change in substrate preference of the PRC2 complex. P527H and C539Y *EZH2^MUT^* retained the wildtype preference of mono- methylated peptide substrate over unmethylated or dimethylated peptides (Figure 7B, C).

As P527H and C539Y *EZH2^MUT^* displayed some enzymatic activity, we investigated if EPZ-6438 would inhibit this remaining enzymatic activity. Despite lower activities of these *EZH2^MUT^*- containing PRC2, there was no change in IC_50_ of EPZ-6438 for either P527H and C539Y *EZH2^MUT^*or *EZH2^WT^* (Figure 7D). We were able to reproduce previously published preclinical and clinical data that EPZ-6438 is a ∼50x more potent inhibitor of EZH2 than EZH1 (13,36). Overall, the enzymatic *in vitro* assessment indicated that the majority of EZH2 mutations resulted in loss of function to various degrees but no change in substrate preference.

## IV. Discussion

Our study indicates several outcomes of EZH2 LOF mutations in disrupting PRC2 activity and rendering the mutant cells dependent on low-level, residual enzyme activity. In the chemical probe screen of epigenetic regulatory proteins, the AML patient cells that harbored EZH2 mutations were most responsive to EZH2 inhibition. This was consistent with *EZH2^MUT^* cells responding to EPZ-6438 by upregulating gene expression signatures enriched in apoptosis and growth arrest. We also note that *EZH2^MUT^*cells had significantly reduced H3K27me3; most had lower levels of EZH2 protein and lower levels of PRC2 components EED and SUZ12. In cases where the *EZH2^MUT^* cells maintained EZH1 protein levels, the reduction of essential complex component EED was consistent with drastically suppressed H3K27me3. However, in keeping with the intrinsically lower catalytic activity of EZH1 (37), the apparent compensatory elevation EZH1 was insufficient to fully restore H3K27me3 levels in EZH2 LOF mutant cells.

Our study identified novel and previously described mutations of EZH2 in MDS/MPN/AML patients. These mutations resided in the SET and CXC domains necessary for the catalytic activity. EZH2-R690H mutation was previously reported in different myeloid disorders such as MDS/MPN (38,39) and leukemia (40). This mutation is well-documented to cause loss of PRC2 function and contributes to chemotherapy resistance (23). Mutation Q653E was reported previously in refractory T-cell leukemia, and V679M was reported in AML, CML, MDS, and myelofibrosis (19,41). T683I was reported in MDS (42), and C539 mutations were reported in MDS (20). Mutations at position P527 were not reported previously in hematopoietic or lymphoid malignancies (COSMIC Server accessed on 06/06/2023). A previous study by Rinke et al also noted multiple *EZH2* mutations in a single patient (18). In our study, the cases with *EZH2^MUT^* allele frequency of >90% did not show gross chromosome 7 aberrations, indicating that other mechanisms, such as microdeletions or loss of heterozygosity, were responsible.

PRC2 is essential in regulating genes responsible for cell differentiation and stemness. Gain and loss of function mutations in EZH2 result in transcriptome changes driving disease (9,18,43-45). Although we noted that EZH2 inhibition resulted in the regulation of several previously reported genes and gene categories and, in general, *EZH2^WT^* cells tended towards gene upregulation while *EZH2^MUT^* patient cells prominently downregulated survival-associated genes, importantly, patient cells showed vastly different baseline transcriptome reflecting their heterogeneous genetic alterations. Heterogeneity in transcriptomes and response to EZH2 knockdown were also noted before (18).

Significant progress has been made in the discovery of PRC2 inhibitors, leading to the approval of tazemetostat as a clinical treatment option (13); furthermore, clinical development of EED inhibitors as allosteric inhibitors of PRC2 is ongoing (46). Together with epigenetic chemical probe tools, these compounds also enabled us to interrogate the vulnerabilities of leukemia cells and identify AML patient cells with sensitivity to PRC2 inhibition. In a leukemia patient cell screen, we found that cells most sensitive to EZH2 inhibitor chemical probe UNC1999 harbored EZH2 mutations. These results were further expanded into additional *EZH2^MUT^* samples and validated using the clinical compound, EPZ-6438, and an EED inhibitor A395. The growth suppression elicited by an EED inhibitor was more prominent, which may be consistent with both EZH2 and EZH1 catalytic function requiring EED, while EPZ-6438 displayed weaker inhibitory action on EZH1. However, both pharmacological PRC2 modulators selectively suppressed growth in EZH2 mutant cells. This selective suppression aligns with a study examining genome-wide copy number and loss-of-function data in a panel of cancer cell lines, demonstrating that partial copy number losses of specific genes rendered cells highly dependent on the remaining copy (47). Furthermore, mutations or copy loss in splicing factors such as SF3B1 can sensitize cells to the knockdown of SF3B1 or spliceosome modulatory drugs (48,49). These observations are in line with the partial loss of tumor suppressor genes leading to carcinogenesis, while the complete loss of these genes was not tolerated. EZH2 plays a dual role acting as a tumor suppressor during AML induction and as an oncogene in disease maintenance (9,10). Thus it is possible that cancer evolution of EZH2 LOF initiates MDS/AML, but also creates an evolutionary trap as the activity of the PRC2 complex is needed in leukemia maintenance and leukemic cell survival.

The *EZH2^MUT^* patient cells had drastically reduced levels of H3K27me3 and lower EZH2 levels but maintained levels of EZH1. Similar findings of EZH2 mutations associated with loss of EZH2 protein were reported in MDS/AML, where coincidentally low EZH2 protein and H3K27me3 levels showed a stronger correlation with reduced patient survival than the presence and content of EZH2 mutation (24,50,51). However, illustrating the complexity, *EZH2^MUT^* patient 177391 cells had higher EZH2 Q653K protein levels and maintained EZH1 and EED levels, which still resulted in undetectable H3K27me3. The increased Q653K EZH2 protein levels could be due to elevated mutant protein stability. On the other hand, 166315 sample cells with E137X nonsense mutation had no detectable EZH2 protein, elevated EZH1, and stable levels of EED and SUZ12. The 166315 cells also maintained low H3K27me3 levels due to EZH1 activity, inhibition of which likely conferred sensitivity to EPZ-6438 and A395. EZH1 has been previously shown to compensate for EZH2 loss (10,52). However, EZH1 levels were not uniformly elevated in *EZH2^MUT^* patient cells. In fact, in another category of *EZH2* mutations (three out of five samples), despite similar mRNA levels, mutant EZH2 protein was plausibly destabilized by mutations, coinciding with reduced protein levels of EED, lower SUZ12 (two out of five), indicating a general disruption of PRC2 complex. Such PRC2 destabilization was reported in human embryonic stem cells upon the knockout of EZH2 (53).

EED inhibitors or the new dual EZH2/1 inhibitors are currently being explored as cancer therapeutics (46). The recent discoveries of mutant histones in cancers indicate that H3K27M mutations in glioma can rewire the chromatin landscape with the remaining PRC2 activity concentrating on the oncogenic loci (54), driving growth suppression programs similar to the regulation we identified in gene expression analysis. Studies also show that the decrease in EZH2 protein levels, especially in relapsed AML, correlates with poor survival in AML (23,24,50). Reduced PRC2 signaling has also been exploited in targeting SMARCA2 and SMARCA4 signaling (55). Loss of

## Notes

### Competing Interest Statement

The authors have declared no competing interest.

